# Membrane curvature and the Tol-Pal complex determine polar localization of the chemoreceptor Tar in *E. coli*

**DOI:** 10.1101/212670

**Authors:** Terrens N. V. Saaki, Henrik Strahl, Leendert W. Hamoen

**Author notes:** For correspondence: L. W. Hamoen.

## Abstract

Chemoreceptors are localized at the cell poles of *Escherichia coli* and other rod-shaped bacteria. Over the years different mechanisms have been put forward to explain this polar localization; from stochastic clustering, membrane curvature driven localization, interactions with the Tol-Pal complex, to nucleoid exclusion. To evaluate these mechanisms, we monitored the cellular localization of the aspartate chemoreceptor Tar in different deletion mutants. We did not find any indication for either stochastic cluster formation or nucleoid exclusion. However, the presence of a functional Tol-Pal complex appeared to be essential to retain Tar at cell poles. This finding also implies that the curvature of cell poles does not attract chemoreceptor complexes. Interestingly, Tar still accumulated at midcell in *tol* and in *pal* deletion mutants. In these mutants, the protein appears to gather at the base of division septa, a region characterised by strong membrane curvature. Chemoreceptors, like Tar, form trimer-of-dimers that bend the cell membrane due to a rigid tripod structure with an estimated curvature of approximately 37 nm. This curvature approaches the curvature of the cell membrane generated during cell division, and localization of chemoreceptor tripods at curved membrane areas is therefore energetically favourable as it lowers membrane tension. Indeed, when we introduced mutations in Tar that abolish the rigid tripod structure, the protein was no longer able to accumulate at midcell or cell poles. These findings favour a model where chemoreceptor localization in *E. coli* is driven by strong membrane curvature and association with the Tol-Pal complex.

**Importance:** Bacteria have exquisite mechanisms to sense and to adapt to the environment they live in. One such mechanism involves the chemotaxis signal transduction pathway, in which chemoreceptors specifically bind certain attracting or repelling molecules and transduce the signals to the cell. In different rod-shaped bacteria, these chemoreceptors localize specifically to cell poles. Here, we examined the polar localization of the aspartate chemoreceptor Tar in *E. coli*, and found that membrane curvature at cell division sites and interaction with the Tal-pol protein complex, localize Tar at cell division sites, the future cell poles. This study shows how membrane curvature can guide localization of proteins in a cell.

## Introduction

Bacteria use specific chemotaxis systems to sense chemical changes in their environment and respond accordingly. One of the best-known systems is that of *Escherichia coli*, which comprises five different membrane spanning chemoreceptors. The cytoplasmic domains of chemoreceptors associate with the adaptor protein CheW and with the histidine kinase CheA. When a receptor binds a specific ligand, CheA is activated and will subsequently phosphorylate the response regulator CheY, which acts on the flagellar motor to change rotation direction. Sensitivity of the chemoreceptors is tuned by methylation and demethylation for which the methylesterase CheB and methyltransferase CheR are responsible. The chemoreceptors are therefore also referred to as methyl-accepting chemotaxis proteins or MCPs. For and in-depth review on the chemotaxis system see e.g. (1, 2).

MCPs form large protein clusters together with CheW, -Y, -A, -B and –R at the cell poles of different bacteria including the Gram-negative model system *E. coli* and the Gram-positive model system *Bacillus subtilis* (3, 4). Several mechanisms have been proposed for this polar localization. In long filamentous *E. coli* cells, YFP labelled CheR clusters were found to assemble with a certain periodicity along the cell axis that corresponds to the position of future division sites. This model is referred to as the ‘stochastic nucleation model’ (5-7). Another theory postulated that MCPs preferably assemble at the curved membrane of cell poles (8). Chemoreceptors form membrane spanning trimers-of-dimers that interact at their cytoplasmic domain at a slight angle thereby forming a tripod-like configuration (9). Consequently, the trimer of dimers prefer bend membrane areas due to the reduced curvature mismatch. (10, 11). This model was recently supported by mechanically bending of whole *E. coli* cells in curved micro-chambers (12), and was also shown to be the main mechanism by which the chemoreceptor TlpA of *B. subtilis* is localized (13). However, another study suggested that polar curvature is not crucial for the localization of chemoreceptor proteins in *E. coli*, but that this requires interaction with the Tol-Pal complex (14). The trans-envelope Tol-Pal complex is a widely conserved component of the cell envelope of Gram-negative bacteria, and is involved in several processes among which cell division (15, 16). In contrast to this, another recent study showed that, at least for the serine chemoreceptor Tsr, the Tol-Pal complex is not required for polar localization, and that nucleoid exclusion is the driving force for polar localization of MCPs (17). Here, we evaluated the different polar localization models in *E. coli* using the aspartate chemoreceptor Tar. We found neither evidence for periodic clustering nor for nucleoid exclusion, but both membrane curvature and the Tol-Pal system appeared to be required for polar localization of Tar in *E. coli*.

## Results

### Stochastic nucleation

The stochastic nucleation model has been based on the formation of large YFP-CheR and CheY-YFP clusters that were regularly spaced with a periodicity of approximately 1 µm (5). To confirm that MCPs also produce these regular clusters, the *E. coli* chemoreceptor Tar was C-terminally fused with a monomeric GFP variant (mGFP). Since GFP tends to form weak dimers, monomeric GFP was chosen to prevent possible localization artefacts (18). To reduce potential artefacts related to protein overexpression, a low copy plasmid with a weakened IPTG-inducible promoter (pTRC99A (19)) was used to express the fusion protein. As shown in Fig. 1A, Tar-mGFP shows a classical septal and polar localization pattern. This localization does not depend on interaction with other chemoreceptors, since expression of the fusion protein in a MCP deletion strain shows the same localization pattern (Fig. S1).

**Fig. 1.**
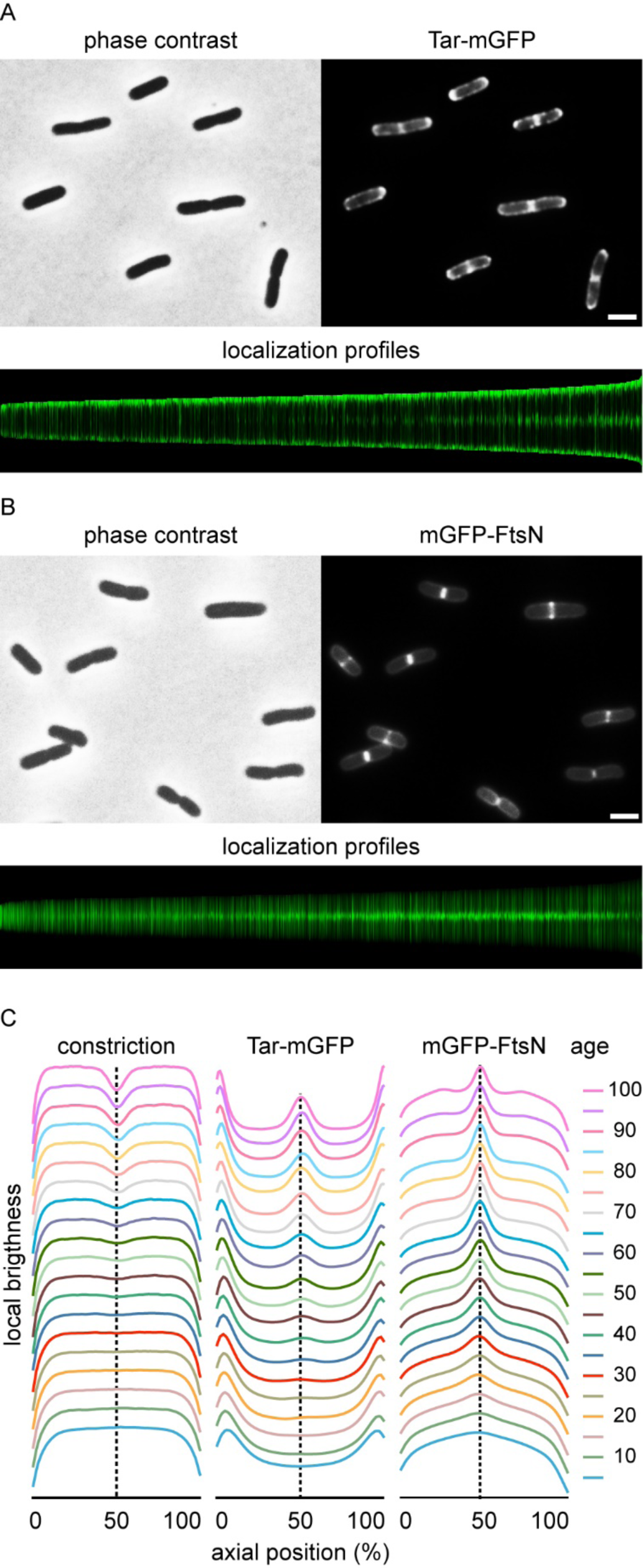
Localization of Tar-mGFP. (A) Fluorescence microscopy image of wild type *E. coli* cells expressing Tar-mGFP. Lower panel shows sorted axial fluorescence profiles, indicative for Tar-mGFP localization during the cell cycle. (B) Fluorescence microscopy image and cell cycle localization profile of cells expressing mGFP-FtsN. (C) Graphical presentation of cell constriction and fluorescence signals during the cell cycle calculated from the localization profiles. Cell age is expressed as % of the cell cycle. 5074 and 6437 cells were used to construct the cell cycle localization profiles for mGFP-FtsN and Tar-mGFP, respectively. Scale bars are 2 µm. Used strains in A and B are TSE29 and TSE48, respectively.

According to the stochastic nucleation model, chemotaxis proteins form large protein clusters prior to the initiation of cell division. To determine at what time in the cell cycle Tar accumulates at midcell, we performed a virtual time lapse approach by sorting cells on size (Fig. 1A, lower panel), and plotting the related fluorescence intensities and cell constriction (Fig. 1C) (20). As a timer for cell division, we followed the localization of GFP-labelled FtsN, an essential cell division protein and part of the cell division machinery (21) (Fig. 1B). Comparison of the localization profiles indicates that Tar appears later at midcell compared to FtsN, suggesting that clustering of the chemotaxis proteins does not precede cell division.

To determine whether Tar forms regularly spaced clusters with a periodicity of around 1 µm in filamentous non-dividing cells, we blocked cell division using the antibiotic cephalexin, which inactivates the cell division protein FtsI required for septum synthesis (22). As shown in Fig. 2A & B, no large regularly spaced fluorescent clusters were observed along the lateral wall of filamentous cells, but the polar clustering remained. Based on these data it seems unlikely that Tar uses stochastic clustering to accumulate at cell poles.

**Fig. 2.**
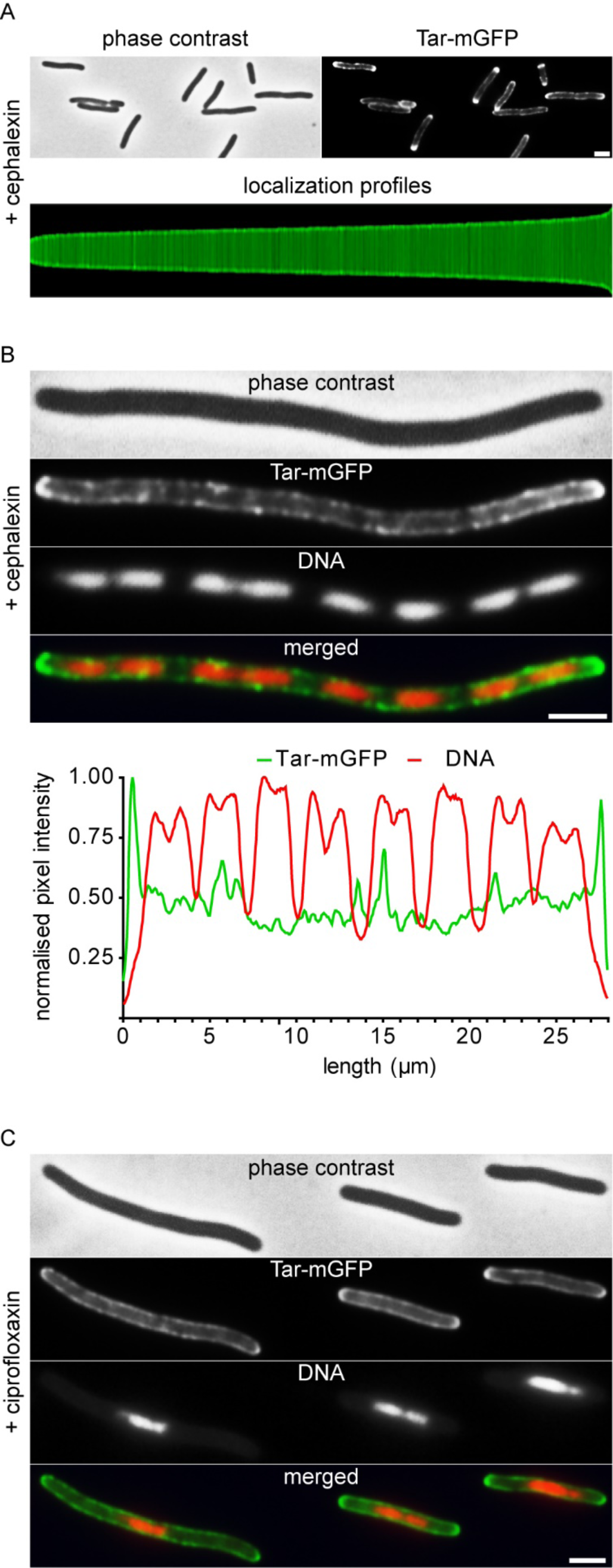
Tar-mGFP clustering and relation to nucleoid position. (A) Fluorescence microscopy image and localization profile of Tar-mGFP expressing cells treated with 15 µg/ml cephalexin for 1 h. 1872 cells were used to construct the localization profile. (B) Fluorescence microscopy image of Tar-mGFP expressing cells treated with 15 µg/ml cephalexin for 3 h. Nucleoids were stained with DAPI. Line scans for the GFP and DAPI signals are presented below. More examples are shown in Fig. S2. (C) Fluorescence microscopy image of Tar-mGFP expressing cells treated with 0.035 µg/ml ciprofloxacin for 1 h. Nucleoids were stained with DAPI. Scale bars are 2 µm. Used strain is TSE29.

### Nucleoid exclusion

In a recent report it was suggested that the serine MCP Tsr of *E. coli* is driven to cell poles by the ‘volume exclusion’ effect of the nucleoid (17). To examine whether nucleoids influence the distribution of Tar, we stained the cephalexin treated cells with the fluorescence DNA dye DAPI to visualize the nucleoids (Fig. 2B). We could not detect any correlation with the position of the nucleoids and the density of Tar-mGFP clusters along the lateral wall (see also line scans in Fig. 2B, and Fig. S2). To corroborate this, cells were treated with ciprofloxacin, which inhibits DNA gyrase and blocks DNA replication, resulting in a dense nucleoid at the centre of long cell, since the activated SOS-response also inhibit cell division (Fig. 2C) (23, 24). Also under these conditions the Tar-mGFP signal was not reduced at the area occupied by the nucleoid. Thus, at least for Tar, nucleoid exclusion does not seem to be important for polar localization.

### CheA stimulated clustering

Clustering of the chemotaxis complex is stimulated by dimerization of the kinase CheA, which interacts with the cytoplasmic domains of the MCPs (25). In fact, we have found that CheA is essential to maintain polar localization of the chemoreceptor TlpA in *B. subtilis* (13). However, it has been shown some time ago that in *E. coli* CheA is not necessary for the polar clustering of chemoreceptor (26). Indeed, when we expressed Tar-mGFP in a *cheA* deletion mutant background, the protein accumulated at midcell and cell poles (Fig. 3A).

**Fig. 3.**
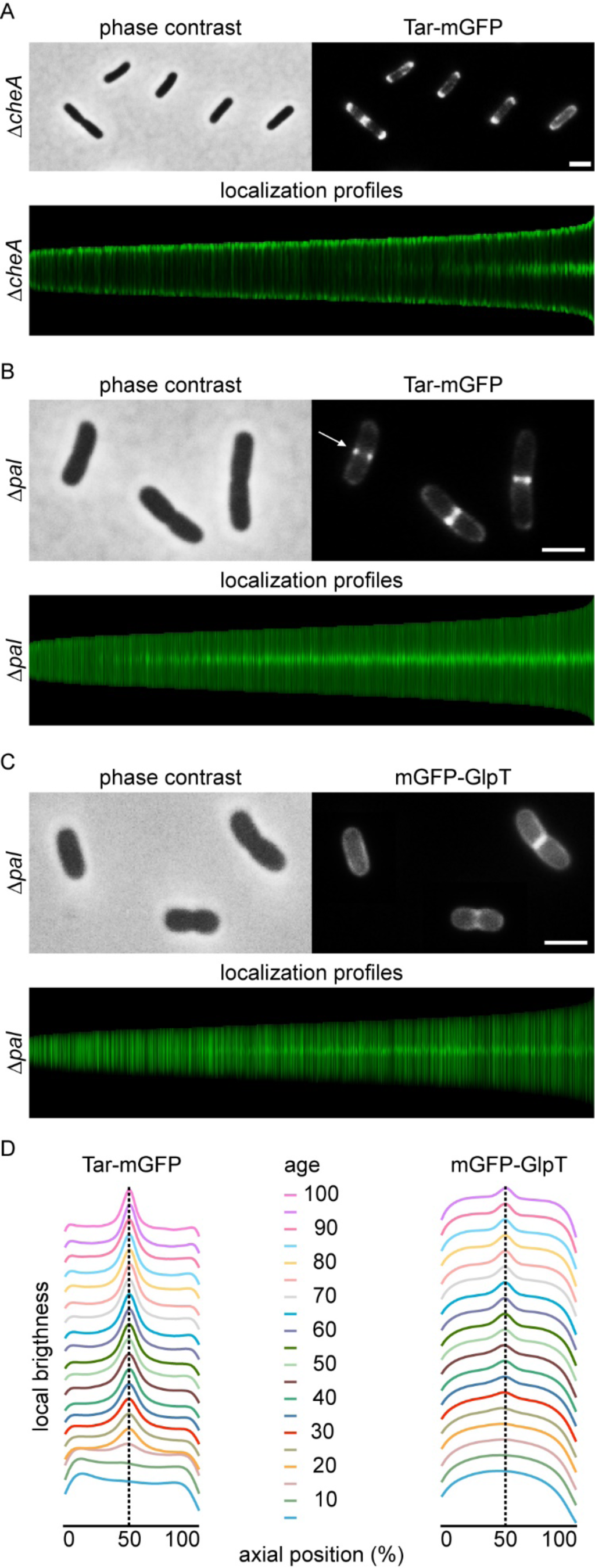
Effect of *cheA* and *pal* deletion mutant on Tar-mGFP localization. (A) Fluorescence microscopy image and localization profile of Tar-mGFP expressing *cheA* deletion mutant. 5948 cells were used to construct the localization profile. (B) Fluorescence microscopy image and localization profile of Tar-mGFP expressing *pal* deletion mutant. Arrow indicates Tar-mGFP foci. 9044 cells were used to construct the localization profile. (C) Fluorescence microscopy image and localization profile of mGFP-GlpT expressed in the *pal* deletion mutant. 5214 cells were used to construct the localization profile. (D) Graphical presentation of fluorescence signals during the cell cycle calculated from the localization profiles. Cell age is expressed as % of the cell cycle. Scale bars are 2 µm. Used strains in A, B and C are TSE38, TSE31 and TSE71, respectively.

### Role of Tol-Pal

Another protein that has been implicated in the polar localization of chemoreceptor proteins is the trans-envelope Tol-Pal complex (14), which accumulates at midcell and assist in the division of the outer cell membrane (16). Pulldown experiments have suggested a direct interaction between TolA and chemoreceptors. However, a recent study questioned the role of the Tol-Pal complex in chemoreceptor localization (17). To verify this, we expressed the Tar-mGFP fusion in a *pal* deletion mutant. Indeed, the polar accumulation of the fusion protein was completely abolished, however, there was still a strong accumulation at midcell, comparable to what is observed in wild type cells (Fig. 3B & D). When the fusion protein was expressed in a *tolA* deletion mutant, a similar localization pattern was observed (Fig. S3). This seemingly contradictory finding (no polar but still midcell accumulation) might explain the different reports on the role of Tol-Pal.

During cell division, a double cell membrane is formed when the division septum is synthesized. This will give a higher fluorescent membrane signal at midcell, so even when Tar is unable to localize and diffuses freely throughout the cell membrane, the extra cell membranes at the division sites could, in theory, account for an increase in GFP signal at midcell. To assess this, we followed the localization of a general transmembrane protein, the glycerol-3-phosphate transporter GlpT (27), throughout the cell cycle in the *pal* mutant. Indeed, the GFP signal showed a slight accumulation at midcell when cells started to divide (Fig. 3 C), however, the signal intensity was much lower compared to that of Tar-mGFP (Fig. 3 D), indicating that the double cell membrane at the division site is not responsible for the strong fluorescence Tar-mGFP signal at midcell.

### Membrane curvature

A closer inspection of the *pal* mutant revealed that the Tar-mGFP signal often appears as two fluorescent dots at midcell (Fig. 3B), suggesting that Tar accumulates as a ring at midcell. The Tol-Pal complex is recruited to the division site by FtsN, and links invagination of the outer membrane with that of the cell membrane during cell division (16). Inactivation of Tol-Pal strongly delays invagination of the outer membrane compared to the cell membrane, and this results in the formation of a division septum that resembles the septal cross walls in Gram-positive bacteria (28). The consequence of such mode of division is that the cell membrane at the transition from the lateral wall to the nascent septal wall is strongly curved (29). This is where the Tar-mGFP fluorescent signal seems to accumulate in the *pal* deletion mutant (Fig. 3B). Presumably, Tar localizes at this region because of the curvature mismatch generated by the tripod configuration of the trimer-of-dimers in combination with the stiffness of the dimers (30). Tension in the membrane is released when these tripods locate to regions of the cell with a corresponding membrane curvature, such as those found at cell division sites. To confirm this, we introduced a N379R mutation in the trimerization site of Tar, corresponding to the N381R mutation in Tsr, which has been shown to abolish trimerization (31). Since single membrane spanning MCP dimers will not deform the membrane, they should therefore not accumulate at cell division sites when membrane curvature is the main driver for localization (Fig. 4A). Indeed, the N379R mutation resulted in the absence of a clear septal and polar fluorescent signal (Fig. 4B & D). When we increased the flexibility of the dimers by introducing a stretch of 3 glycines in the HAMP domain (G248G D249G L250G) of the dimers, Tar was also no longer able to accumulate at midcell and cell poles (Fig. 4C & D), in line with the assumption that the unstructured glycine stretch eliminates the membrane curvature preference of the trimer (Fig. 4A) (12, 13). Thus, membrane curvature seems to drive the localization of Tar trimers-of-dimers.

**Fig. 4.**
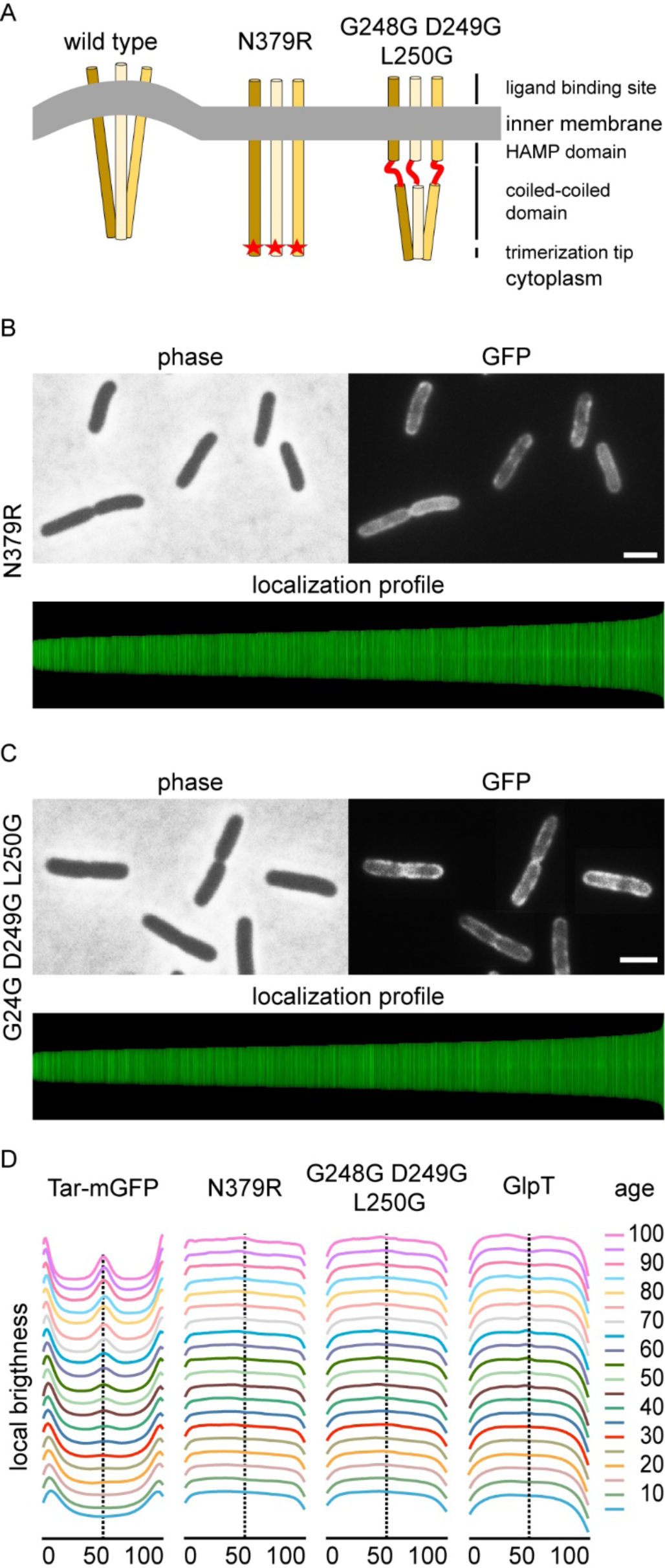
Membrane curvature is important for localization. (A) Schematic presentation of the effect of dimerization mutation N379R and the triple glycine insertion (G248G D249G L250G) on membrane curvature mismatch. (B) Fluorescence microscopy image and localization profile of Tar(N379R)-mGFP expressing cells. 7331 cells were used to construct the localization profile. (C) Fluorescence microscopy image and localization profile of Tar(G248G D249G L2450G)-mGFP expressing cells. 7856 cells were used to construct the localization profile. (D) Graphical presentation of fluorescence signals during the cell cycle calculated from the localization profiles. Cell age is expressed as % of the cell cycle. Scale bars are 2 µm. Used strains in B and C are TSE41 and TSE42, respectively, and in D strains TSE29 and TSE67.

## Discussion

Our data suggests that the MCPs of *E. coli* and *B. subtilis* arrive at cell poles by a comparable mechanism. First, they accumulate at midcell when during cell division the cell membrane takes on a strong concave (seen from the cytoplasmic face) shape at the base of the nascent septum. Based on a composite crystal structure the curvature of one trimer-of-dimers was calculated to amount to a radius of approximately 37 nm (8). The membrane curvature mismatch is reduced considerably when a trimer-of-dimer is located at the base of the nascent division septum. After septation is completed, *B. subtilis* chemoreceptor clusters are maintained at the newly formed poles by forming large protein clusters that require CheA. However, in *E. coli*, the Tol-Pal complex is required to keep chemoreceptors clustered at the newly formed cell poles, instead of CheA. Co-immunoprecipitation experiments have suggested a direct interaction between this complex and chemoreceptor proteins (14). Since lateral diffusion of the large trans-envelope Tol-Pal complex is likely to be hampered by the peptidoglycan layer, interactions between Tol-Pal and chemoreceptor proteins might anchor MCPs and maintain their polar localization.

Several papers have argued that the curvature of the *E. coli* cell pole is sufficient to attract MCP trimers (8, 12, 32). However, the polar localization of Tar-mGFP is completely abolished when Tol-Pal is absent. This indicates that the curvature at the cell pole is not sufficient to markedly reduce the membrane curvature mismatch created by the Tar trimer-of-dimers. This is maybe not so surprising since the cell pole has a curvature with a radius of approximately 500 nm, which is much larger compared to the 37 nm radius of MCP trimer-dimers (8). Moreover, the cylindrical later wall has a radius that is comparable to that of the cell pole, which makes the perceived curvature increase of cell poles even smaller.

Over the years different mechanisms have been postulated and contradictory results have been obtained in the research of polar localization of *E. coli* chemoreceptors. One possible explanation is that different groups use different protein reporters for chemoreceptors. In many studies, the cytoplasmic CheR has been used as a proxy for chemoreceptor clusters (5, 12, 14), while others have looked directly at the localization of MCPs (17). Another reason might be the use of fluorescent protein reporters with a tendency to dimerize, such as YFP and GFP. This characteristic has been shown to cause localization artefacts, especially when used with proteins that form multimers (18). Finally, we cannot exclude that different chemoreceptor use different mechanisms for localization. Nevertheless, both in *E. coli* and in *B. subtilis* it appears that the strong curvature generated during cell division is a key driving force for the localization of MCPs.

## Materials and methods

### Bacterial strains and growth conditions

All strains used in this study are listed in Table S1. Strains were grown in GB1 minimal medium (6.33 g/l K_2_HPO_4_.3H_2_O, 2.95 g/l KH_2_PO_4_, 1.05 g/l (NH_4_)_2_SO_4_, 0.10 g/l MgSO_4_.7H_2_O, 28 mg/l FeSO_4_.7H_2_O, 7.10 mg/l Ca(NO_3_)_2_.4H_2_O) supplemented with vitamin B1 (4 mg/ml) and 0.4 % glucose as carbon-source, as previously described (33, 34). Auxotrophic BW25113 cells required arginine (50 µg/ml), glutamine (50 µg/ml), uracil (20 µg/ml), and thymidine (2 µg/ml). Either 100 µg/ml (or 5 µg/ml in cases of *pal* or *tolA* mutant) of ampicillin or 25 µg/ml of chloramphenicol was added to the growth medium to maintain plasmids.

### Plasmid construction

Purified DNA amplicons were used in a 1:10 molar ratio of vector to insert(s) in Gibson Assembly reaction (20 µl) at 50^o^C for 60 minutes. 5 µl of each Gibson Assembly reaction mix was used to transform ultra-competent *E. coli* TOP 10 cells. Ultra-competent *E. coli* TOP 10 cells were prepared as described in Hanahan *et al.* (35). Plasmids were sequenced to confirm constructs. Plasmids were transformed into chemically competent BW23115 wild type or mutant cells, prepared as described in Maniatis et al. (36). Transformants were selected on selective LB agar plates containing the appropriate concentration of antibiotic. Oligos (Table S3) and plasmids (Table S2) used in this study are listed in the supplementary information.

To construct Tar-mGFP fusion, the point mutation GFP(A206K) was introduced in plasmid pBAD24-Tar-GFP (37) to prevent dimerization of GFP (18). The mutation was made by quick change using primer pair GFP(A206K)-for/GFP(A206K)-rev, resulting in plasmid pBAD24-Tar-mGFP. To express Tar-mGFP from a weakened isopropyl ß-D-thiogalactoside (IPTG)-inducible promoter (19) and low copy number plasmid, the pBAD promoter was replaced by the pTRC99A promoter from pSAV57 (33), and the pSC101 origin with origin m pSEN29 (38). First, pBAD24-Tar-mGFP was linearized by PCR amplification using primer pair TerS327/TerS328, then the pSC101 origin was amplified with primer pair TerS425/TerS426, and subsequently both products were ligated by Gibson Assembly (39), resulting in plasmid pTNV107 (pBAD24-Tar-mGFP-pSC101 ori). To obtain the weak IPTG-inducible low copy number plasmid, plasmid pTNV107 was linearized with primer pair TerS425/TerS507, and the pTRC99A promoter was amplified from pSAV057 using primer pair TerS328/TerS506. The products were ligated using Gibson Assembly, resulting in pTNV149 (pTRC99A-Tar-mGFP-pSC101 ori).

To test if curvature caused by trimer-of-dimers is essential for Tar-mGFP localization, we introduced a N379R point mutation in Tar that abolishes the interaction between dimers. The primer sets TerS328/TerS517 and TerS425/457 were used to introduce N379R in pTNV149 (pTRC99A-Tar-mGFP-pSC101 ori) using Gibson Assembly, resulting in plasmid pTNV154 (pTRC99A-Tar(N379R)-mGFP-pSC101 ori). We also introduced a stretch of 3-glycines in the HAMP domain of Tar to make the dimers flexible. Primer pairs TerS328/TerS516 and TerS425/515 were used to introduce G248G D249G L250G in Tar in pTNV149 (pTRC99A-Tar-mGFP-pSC101 ori), resulting in plasmid pTNV153 ((pTRC99A-Tar(G248G D249G L250G-mGFP-pSC101 ori).

To compare midcell localization of Tar-mGFP to divisome assembly, we used the late cell division protein FtsN fused to monomeric GFP. The mGFP-FtsN fusion was constructed by PCR amplification of pTNV149 with primer pair TerS418/520, a monomeric variant of *gfp* from pTNV100 with primer pair TerS362/521, and *ftsN* from *E. coli* genomic DNA with primer pair TerS523/541 followed by Gibson Assembly, resulting in plasmid pTNV155 (pTRC99A-mGFP-FtsN-pSC101 ori).

As a control for membrane localization, we constructed a glycerol-3-phosphate transporter GlpT-GFP fusion. mGFP-GlpT was made by PCR amplification of pTNV149 with primer pair TerS418/520, a monomeric variant of *gfp* from pTNV100 with primer pair TerS362/521, and *glpT* from *E. coli* genomic DNA with primer pair TerS544/545, followed by Gibson Assembly, resulting in plasmid pTNV162 (pTRC99A-mGFP-GlpT-pSC101 ori).

### Microscopy and image analysis

The virtual time lapse is based on the fact that during steady state growth the average mass of all cells and their age frequency distribution are constant allowing precise spatio-temporal information on protein localization during the cell cycle, as described in (40). Steady state was obtained by growing cells in GB1 medium at 30 ^o^C under shaking (210 rpm) while keeping OD_450_ below 0.2 by regular dilution in pre-warmed medium for three days to reach steady state growth. At steady state, Tar-mGFP was induced with 15 µM of IPTG for at least two-doubling times. Steady-state cells were centrifuged at 1000 RPM for 2 minutes to bring the OD_450_ to ~0.4. 0.3 µl cells were spotted onto a microscope slide covered with a thin layer of 1.3% agarose. When applicable, cells were treated with 15 µg/ml cephalexin for 1 to 4 h, or 0.035 µg/ml ciprofloxacin for 1 h. Images were acquired with 500 ms exposure time for the GFP channel. Fluorescent microscopy was carried out with a Nikon Eclipse Ti equipped with a CFI Plan Intensilight HG 130 W lamp, a C11440-22CU Hamamatsu ORCA camera, and NIS elements software, version 4.20.01. Images were analysed using Image J v 1.50i (https://imagej.nih.gov/ij/) and the Image J plugin ObjectJ version 03p (40).

## Acknowledgements

We thank the members of the Bacterial Cell Biology group for useful discussions, especially Tanneke den Blaauwen for discussions on protein localization in *E. coli*. We would like to thank Séverin Ronneau (Imperial College London) for constructing the initial monomeric GFP reporter, and Ikuro Kawagishi (Hosei University), Joen Luirink (VU University), Pierre Genevaux (CNRS), and Tom Shimizu (AMOLF) for kindly providing us with strains and plasmids. This research was funded by a Biotechnology and Biological Sciences Research Council (BBSRC) grant BB/I01327X/1, Marie Curie CIG grant DIVANTI (618452), and NWO STW-Vici grant 12128.

